# Bioptim, a Python framework for Musculoskeletal Optimal Control in Biomechanics

**DOI:** 10.1101/2021.02.27.432868

**Authors:** Benjamin Michaud, François Bailly, Eve Charbonneau, Amedeo Ceglia, Léa Sanchez, Mickael Begon

**Author notes:** These authors have contributed equally to this work and share first authorship (Benjamin Michaud, François Bailly).

## Abstract

Musculoskeletal simulations are useful in biomechanics to investigate the causes of movement disorder, to estimate non-measurable physiological quantities or to study the optimality of human movement. We introduce *Bioptim*, an easy-to-use Python framework for biomechanical optimal control, handling musculoskeletal models. Relying on algorithmic differentiation and the multiple shooting formulation, *Bioptim* interfaces nonlinear solvers to quickly provide dynamically consistent optimal solutions. The software is both computationally efficient (C++ core) and easily customizable, thanks to its Python interface. It allows to quickly define a variety of biomechanical problems such as motion tracking/prediction, muscle-driven simulations, parameters optimization, multiphase problems, etc. It is also intended for real-time applications such as moving horizon estimation and model predictive control. Six contrasting examples are presented, comprising various models, dynamics, objective functions and constraints. They include data-driven simulations (i.e., a multiphase muscle driven gait cycle and an upper-limb real-time moving horizon estimation of muscle forces) and predictive simulations (i.e., a muscle-driven pointing task, a twisting somersault with a quaternion-based model, a position controller using external forces, and a multiphase torque-driven maximum-height jump motion).

## I. Introduction

Biomechanics researchers rely on numerical simulations of motion to gain understanding on a variety of scientific topics such as the physiological causes of movement disorders and their consequences on health [1], the estimation of non-measurable physiological quantities (e.g., muscle forces [2]) and the optimality of human movement [3]. The musculoskeletal models used in these simulations generally have a large number of degrees of freedom and they are governed by several ordinary differential equations (ODEs) which mainly describe multibody and muscle activation dynamics. The complexity of these systems has led scientists to formulate their simulations as optimal control problems (OCP), relying on efficient non-linear optimization software to find trajectories that fulfill a desired task while enforcing the system dynamics and minimizing a cost (e.g. motion duration, energy expenditure, matching experimental data, etc.). Up to very recently, there was no off-the-shelf software available to the community to quickly formulate and solve such musculoskeletal OCPs [4]. Consequently, researchers had to develop their own solutions, with little or no dissemination to the community, limiting synergies between researchers. As a result, many approaches coexist to formulate and solve OCPs in the biomechanical literature. The formulation, also called discretization, consists in turning a continuous trajectory optimization problem into a generic discrete non-linear program (NLP) that is solved using a dedicated algorithm. The main family of so-called *direct* transcription methods comes from numerical optimal control. They consist in straightforwardly choosing the state and/or the control as optimization variables at a given number of points along the trajectory and they rely on the integration of the system dynamics between these points.

For instance, the *direct collocation* method has shown its efficiency in some studies investigating human motion [5], [6]. It consists in approximating the integration of the system dynamics using polynomials that describe the state and control trajectories. Its main advantages are that it leads to very sparse NLPs, that knowledge about the state trajectory can be used in the initialization, and that it handles unstable systems well. Its major disadvantage is that adaptive integration error control implies regridding the whole problem and thus changes the NLP dimensions [7]. *Direct multiple shooting* is another direct method that was also applied with success in a lot of biomechanics [8], [9], [10], [11] and robotics [7], [12], [13] studies. Its advantages are mostly the same as for direct collocation in addition to combining integration error control with fixed NLP dimensions, as it relies on possibly adaptive ODE solvers to integrate the system dynamics. Besides direct methods, other choices can be made, as in [14], [15], where the optimization variables are instants at which a switch in the motor strategy occurs, using polynomials function (4th, 5th order) in-between, or in [16], [17], where the optimization variables are the coefficients of fourth order polynomial approximations of the states, with linking conditions to enforce the continuity of the controls. These last approaches are less generic than the direct methods as they either require a prior knowledge about the state and control trajectories. Most of the time, when investigating complex biomechanics issues, we do not have this information.

Concerning the non-linear solver, a variety of software exist and have been used to solve transcribed musculoskeletal NLPs. They can use different heuristics: interior point methods (*Ipopt*, [18]) or sequential quadratic programming (*snopt* [19], *ACADOS* [20]), but they are all gradient based. Therefore, derivatives of the NLP cost function and constraints are required to perform optimization. These derivatives can be obtained by finite differences (often implemented but inaccurate thus comprising convergence) or computed exactly using automatic differentiation (requiring to write all dependencies of the software in symbolic variables), using, e.g., *CasADi* [21].

In order to promote the use of musculoskeletal optimal control among biomechanics researcher, we identified a strong need for a dedicated tool, as shown by the recently launched *OpenSim Moco* [22]. The biomechanics community being mainly composed of software users, such a tool should request a flexible user interface written in a widely used high-level and if possible open-source language (e.g. Python) with a low-level core (e.g. C++) for efficiency. To develop such a software, four interrelated components are essential in our opinion: *i)* a musculoskeletal modeling software, with a visualization module (multibody kinematics and dynamics, muscle dynamics, etc.), *ii)* a method for automatic differentiation, *iii)* a discretization approach, and *iv)* one or several nonlinear programming (NLP) solvers. General-purpose optimal control software (e.g. *GPOPS-II* [23], *Muscod-II* [24], [25], *Acado* [26]]) address *ii)* to *iv)* but they need to be interfaced with a musculoskeletal modeling module and they do not provide any built-in biomechanics features (physiological cost functions, kinematic constraints, etc.). In that sense, the aforementioned *OpenSim Moco*, is a welcome initiative that draws its strength from its integration with the widely used *OpenSim*. However, it faces the following limitations: it uses finite differences to avoid the complexity of adapting the *OpenSim* codebase to support automatic differentiation, it uses direct collocation as transcription method, preventing the use of adaptive ODE solvers and it is not as flexible as required by the community, since it requires the user to develop new features, such as new objective functions, in C++.

The objective of the present paper is to introduce *Bioptim*^1^, *an open source optimal control software dedicated to musculoskeletal biomechanics. Bioptim* is based on C++ code for computational efficiency but the user interface is written in Python for flexibility and ease-of-use. The OCP transcription uses direct multiple shooting to preserve the possibility of using arbitrarily accurate ODE solvers for the integration, which is fully parallelized for more efficiency. *Bioptim*’s core is fully written in *CasADi* symbolics to benefit from algorithmic differentiation and to exploit *CasADi*’s interface with several non-linear solvers (*Ipopt, SNOPT*). Moreover, *Bioptim* is interfaced with the cutting-edge solver *ACADOS*, a recent NLP solver dedicated to direct multiple shooting, intended for real-time applications. The purpose of *Bioptim* is to allow fast and flexible musculoskeletal OCP formulation and solving by providing a framework with a lot of typical biomechanics problem already implemented and customizable.

The paper is organized as follows: first, the design and implementation of *Bioptim* are described. Next, the versatility and performances of *Bioptim* are shown through a variety of examples available online.

## II. Implementation and Design

### A. Implementation and dependencies

*Bioptim* is the top layer of a series of libraries (*Biorbd*: dynamics and MSK modeling; *CasADi*: automatic differentiation; *Ipopt*/*ACADOS*: optimization; *Bioviz*: visualization). Within this software suite, *Bioptim*’s main role is to shape the problem to allow its dependencies to communicate efficiently, while providing an intuitive and flexible interface to the user (Fig. 1). Therefore, it was written in Python for its flexibility and its widespread use among researchers. However, all intensive calculations behind the interface are performed in C/C++, keeping *Bioptim* both fast and easy to customize.

**Fig. 1:**
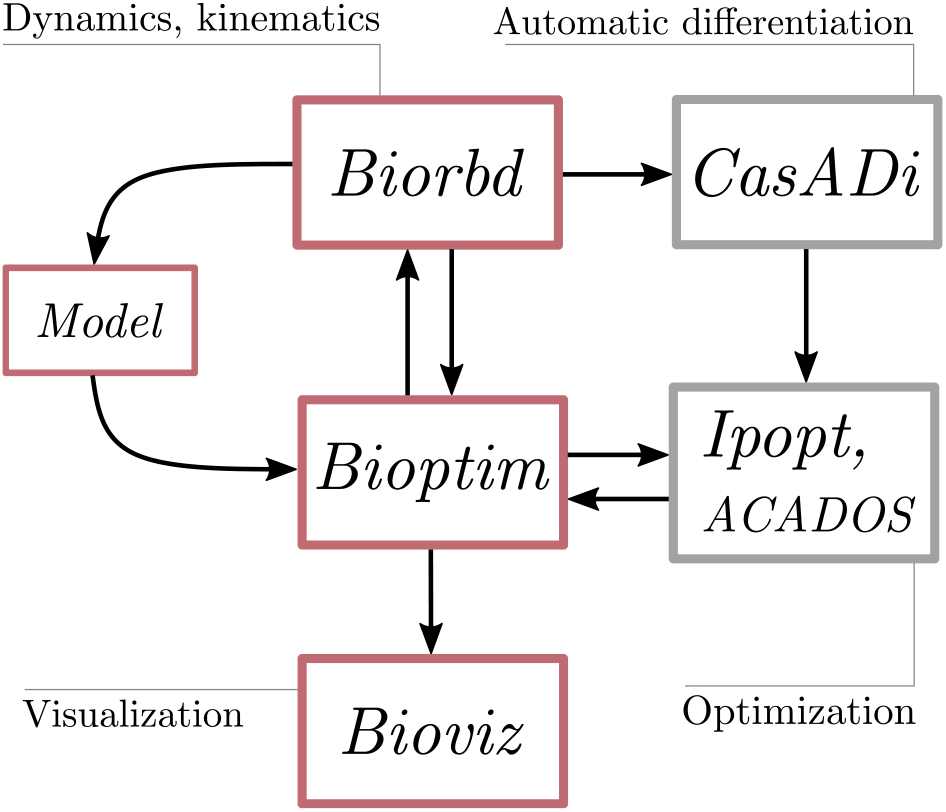
*Bioptim* dependencies flowchart. The red-boxed software are developed by the S2M team. The *Bioptim* part is further detailed in Fig. 2.

**Fig. 2:**
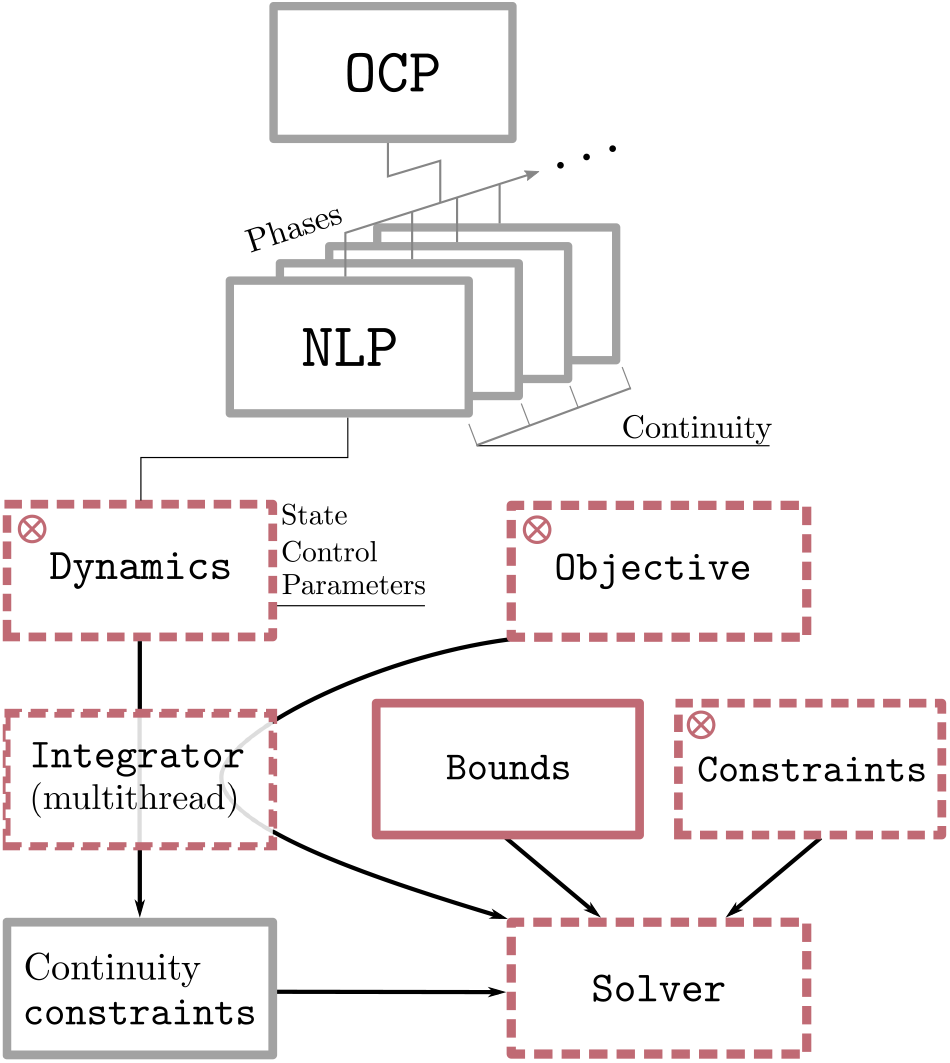
*Bioptim* design flowchart. Red boxes correspond to objects that must be filled in by the user. Red-dashed boxes correspond to pre-implemented objects already available to the users. ⊗ stands for easily customizable objects.

### B. Design

*Bioptim* shapes and solves optimal control problems whose two required entries are a model (.*bioMod* file) and an OCP. The model file contains the geometrical characteristics and the segment inertial parameters as well as optional elements, namely, the markers, the actuators of the model (muscles and joint torques possibly with torque/angle/velocity relationships) as well as bounds on joint kinematics and torques. It also allows the user to design or import meshes for visualization purposes. The `OCP` consists in a combination of nonlinear problems (NLPs) that allows for the formulation of multi-staged OCPs. Each `NLP` has the following attributes: *1)* a dynamics type, *2)* an objective function set, *3)* a constraint set, *4)* variables bounds, *5)* a number of shooting points and the duration of the problem and *6)* initial guesses. Based on these inputs, *Bioptim* properly sets up the multiple shooting transcription of the OCP, with appropriate continuity constraints (between the shooting nodes and the phases) and shapes it up to feed the chosen nonlinear solver (*Ipopt* or *ACADOS*). Next, we develop the different attributes of each `NLP`:

#### 1) Dynamics

The dynamics defines which variables are states (**x**), controls (**u**) and parameters (**p**), the latter being time-independent. Then, it implements the ordinary differential equation governing the state dynamics:

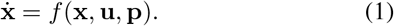

More than 10 dynamics are already implemented in *Bioptim*, among which the controls (piecewise constant or linear) can be muscle excitations, muscle activations and/or joint torques, the states can be muscle activations and/or joint kinematics. They can include contact points, external forces, etc. Even if these dynamics types exhaustively span the current usages in biomechanics, a custom dynamics type is also pre-implemented to easily customize problems.

#### 2) Objective function set

In line with the optimal control formalism, there are two main types of objective functions, namely Lagrange and Mayer. Lagrange types are running objectives, integrated over the NLP duration. Mayer types are time-specific objectives. Classically, they correspond to a terminal objective, but to be more versatile, they can be defined at any instant in *Bioptim*.

`Objective functions` can depend on any of the optimization variables, *i*.*e*. the controls, the states, the parameters and the duration of the problem. A lot of objective function types are already implemented in *Bioptim* (*>* 20), among which tracking / minimizing, on states / controls / markers / contact forces / problem duration, etc. Should one go missing, a custom objective type is also possible to define.

When declaring the desired list of objective functions for a given NLP, each objective function type is associated with a weight, and the user can choose on which components of the vector variables the objective must apply. If applicable (for tracking objective functions mainly), the user must also specify the numerical target of the objective.

#### 3) Constraint set

Classically, constraints are hard penalties of the optimization problem, i.e., a solution will not be considered optimal, unless all constraints (equality or inequality) are met. The `Constraint` class contains a variety of implemented constraints. Some of them are specific functions, commonly useful in biomechanical problems (e.g. non-slipping contact point, non-linear bounds on torque depending on the state, etc.), the others have their equivalent in the `ObjectiveFunction` class. Should one go missing, a custom constraint type is also possible to define.

#### 4) Bounds

Essentially, the `Bounds` are constraints directly related to the states, the controls and the parameters. They are useful to define model-related constraints such as kinematic, torque or muscle excitation / activation limits.

#### 5) Shooting points and problem duration

In direct multiple shooting, the total duration of the problem is divided into smaller intervals whose initial values are called shooting points. In *Bioptim*, the user is asked to define a number of shooting points and a problem duration, per phase. Possibly, the problem duration can be part of the optimization variables, allowing for, e.g., minimal time formulations.

#### 6) Initial guesses

The user can provide an `InitialGuess` for all the optimization variables, at each shooting point. This feature aims at providing prior information to the solver. Several `InterpolationTypes` are implemented (constant, linear, spline, each point, etc.), to quickly let the user define the initial guesses. A custom `InterpolationType` is also possible to implement.

## III. Examples

In this section, six applications are presented to illustrate the versatility of *Bioptim* and give a practical overview on how to use its main features. The settings and performances (convergence time, single shooting integration error, optimized objective) of each OCP are summarized in Tab. I. When possible, problems were solved with both *Ipopt* and *ACADOS*. In the following, bold symbols denote vectors and starred ones (^*^) denote reference or tracked quantities.

**TABLE I:**
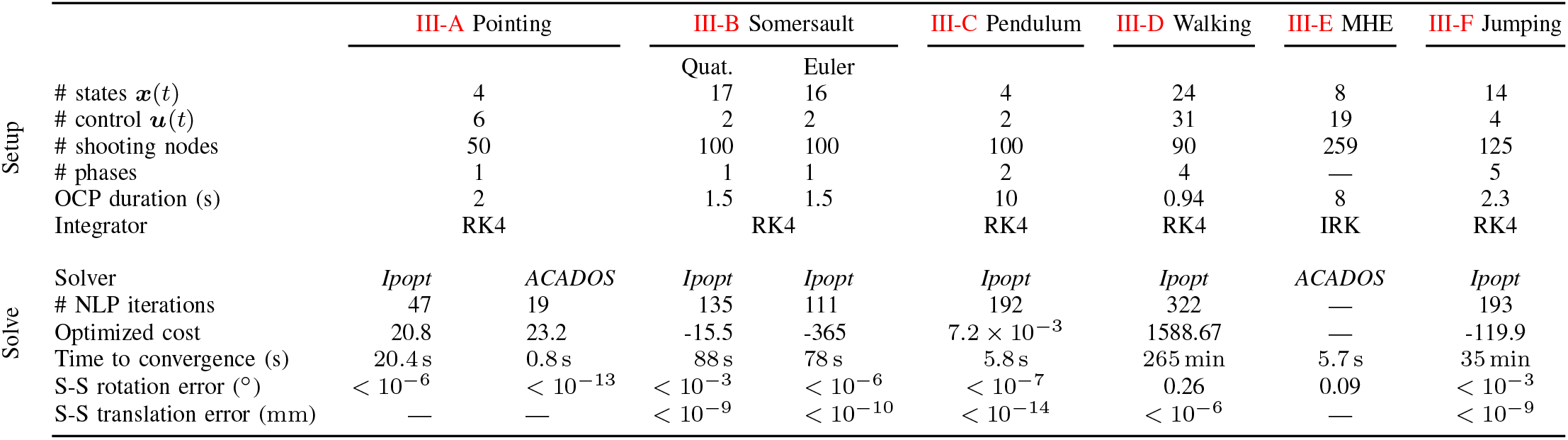
Overview of the computational results for the different examples. The single shooting (S-S) state trajectory is obtained by forwardly integrating the initial state with the optimized control inputs. The S-S error is computed as the average error between the optimized state vector and the single shooting one at *t* = min(1, *T*), with *T* the total duration of the OCP in seconds. Both the rotational and the translational parts of this error are reported in ^*°*^ and mm respectively. — stands for non applicable. All tests were conducted on a personal laptop with an Intel® Core™ i5-8265U CPU @ 1.60GHz × 8, with 24 Gb RAM.

### A. Muscle activation driven pointing task

In this first example, the goal was to achieve a muscle activation driven pointing task using a 2-DoF arm model with six muscle elements. In addition to muscle-induced torques, pure joint torques were added to compensate for the model weaknesses. The main term (highest weight) of the objective function (Eq. 2) is a Mayer objective, corresponding to the pointing tasks at the final node, to superimpose two markers, the first one (**m**_**u**_) fixed in the ulna system of coordinates and the second one 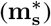 fixed in the scene. The three Lagrange terms were added for control (muscle activation **a** and joint torques **τ**) and state (x) regularization:

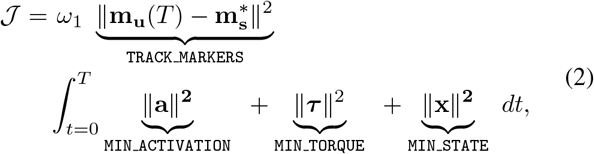

where *T* = 2 s is the duration of the motion, and *ω*_1_ = 1*e*5. The problem was discretized using 50 shooting nodes with a 5-steps Runge-Kutta-4 (RK4) integration in-between. The problem was solved using *Ipopt* (with exact Hessian computations) and *ACADOS* (with a Gauss-Newton approximation of the Hessian) resulting in two very close solutions. *ACADOS* was about 50 times faster than *Ipopt* and was better at enforcing the continuity constraints (as shown by the single shooting error in Tab. I). *Ipopt* however ended up with a smaller optimized objective (20.8 *vs* 23.2), leading to a more optimal solution than *ACADOS*. Superimposed snapshots of the optimal motion found with *ACADOS* are displayed in Fig. 3. It is worth mentioning that for the purpose of this illustration, no constraint was given on the shoulder range of motion to ensure physiological muscle trajectories.

**Fig. 3:**
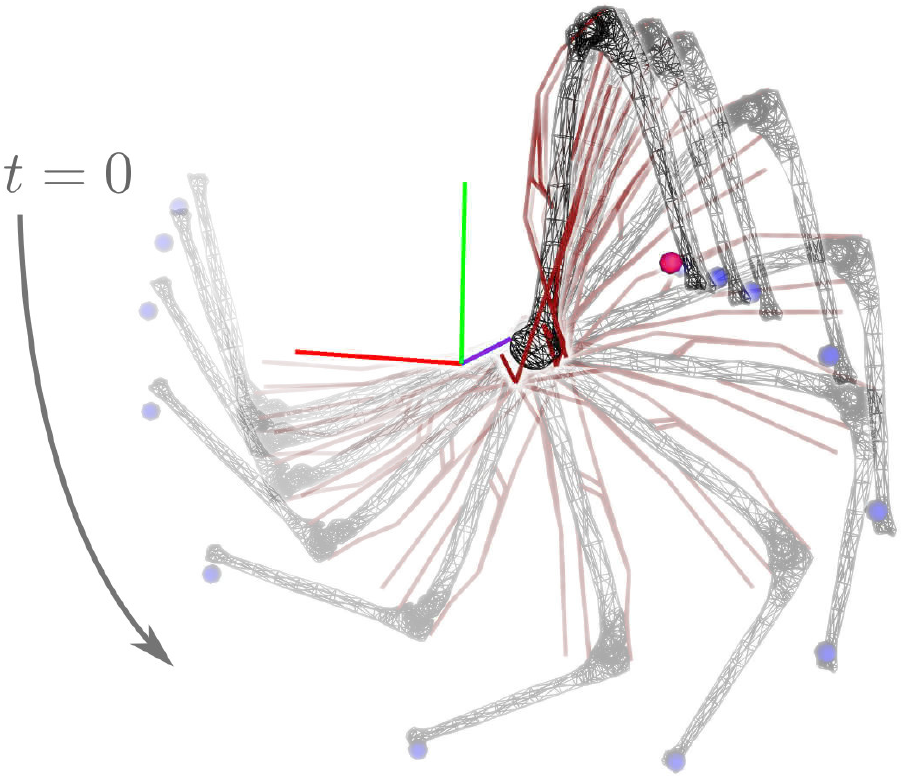
Snapshots of an optimized activation-driven pointing task with *ACADOS*. The arm starts facing upwards in left hand part of the picture and ends facing downwards in the right hand part. The marker fixed on the ulna head is depicted in blue and the scene-fixed target marker is depicted in red. Red lines show the lines of actions of the muscles.

### B. Quaternion base twisting somersault

In this example of acrobatic sports biomechanics, the goal was to maximize the twist rotation (*ϕ*) of an 8-DoF model in a backward somersault. It illustrates *Bioptim*’s ability to handle quaternionic representations of rotations. The model was composed of a 6-DoF root segment and two 1-DoF torque actuated shoulder joints. Two different numerical descriptions of the root segment rotations were used: Euler angles and quaternions. The objective function was as follows:

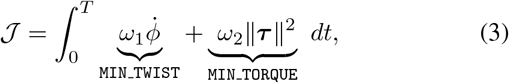

with *ω*_1_ = −1 (resulting in the maximization of the first term) and *ω*_2_ = 10^*−6*^, T is the duration of the movement and ***τ*** is the torque control vector. The first term of the objective function (Eq. 3) corresponds to maximizing the change in twist rotation and the second term is for control regularization.

The movement lasted for approximately 1 second and was discretized with 100 shooting nodes, a kinogram is presented in Fig. 4. The optimal kinematics were different for the two types of models (Fig. 5) because of the presence of local minima. However, both models take profits of a common biomechanical strategy (i.e. tilting the body to bring closer together the twist axis and the angular momentum vector) highlighting the equivalence of the two rotation representations. Euler angles have the advantage to be easily interpretable, but they suffer from the loss of a DoF at the gimbal lock (leading to numerical instabilities). The use of a quaternion-based representation tackles this numerical stability issue when a joint is free to rotate on a three-dimensional range of motion. Quaternion’s integration must be handled with care [11]. Indeed, when representing orientations, quaternions must be unitary and thus belong to a constrained manifold (namely, the unit 3-sphere *S*^3^). However, classical numerical integration schemes such as Runge–Kutta methods treat unit quaternions as if they were arbitrarily defined in R^4^. To overcome this challenge, *Bioptim* performs a normalization after each Runge–Kutta iteration to project non-unitary quaternions onto *S*^3^.

**Fig. 4:**
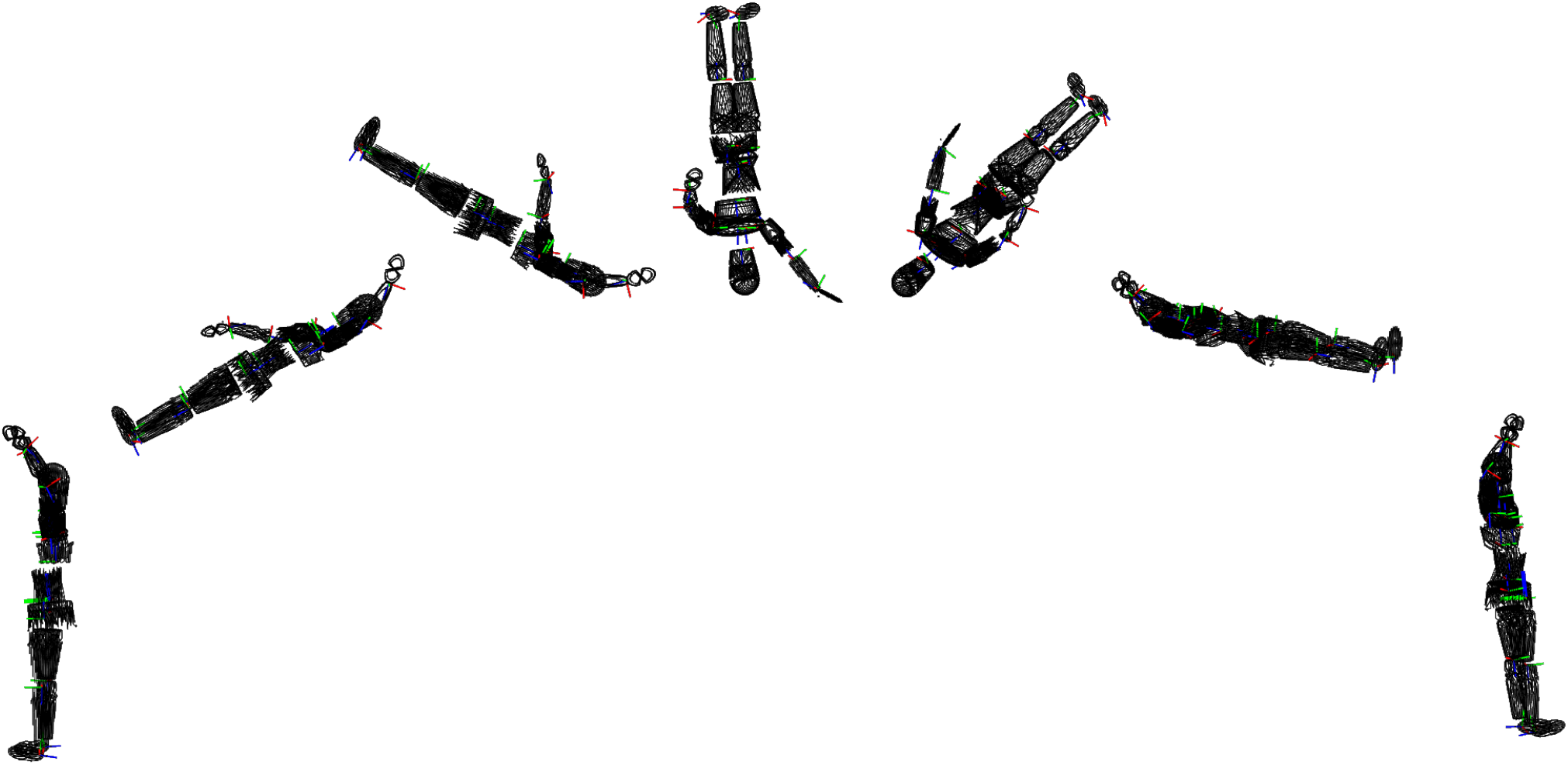
Snapshots of maximally twisting somersaults driven by shoulder torque actuators and a free base whose rotation is either expressed by Euler angles (top) or by quaternions (bottom).

**Fig. 5:**
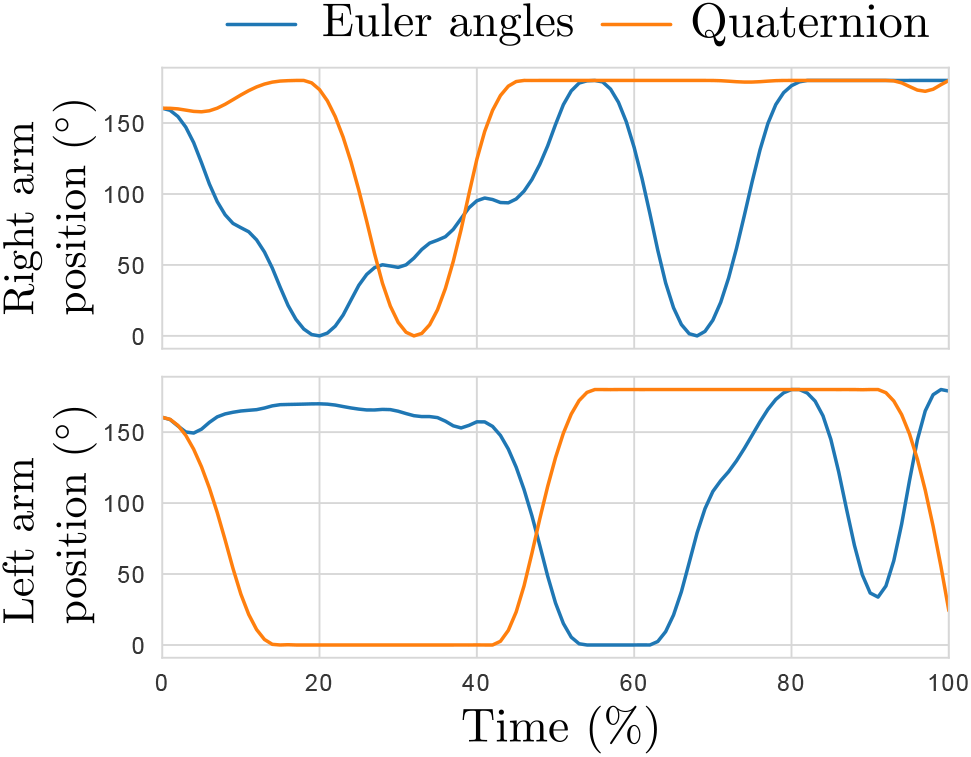
Right (top) and left (bottom) arm kinematics of the twisting avatar for the Euler angles (blues line) and the quaternion (orange line) representation of the orientation of the free base.

### C. Pendulum on a spring

This example is presented to introduce *Bioptim*’s ability to use external forces. The goal was to hold the position of a 1 kg mass hanging on a linear spring attached to the ground. A 0.2 m-long pendulum weighting 10 kg was attached to the mass and free to rotate in one dimension (Fig. 6). In addition to the spring force, the mass was actuated by a vertical external force (e.g., something pulling on it) while the pendulum rotation was passive. The system therefore comprised two DoFs, the mass position (*q*_*m*_) and the pendulum angle (*q*_*p*_) and one control input, the vertical external force pulling on the mass (*τ*). The spring force *ℱ*_*s*_ was:

**Fig. 6:**
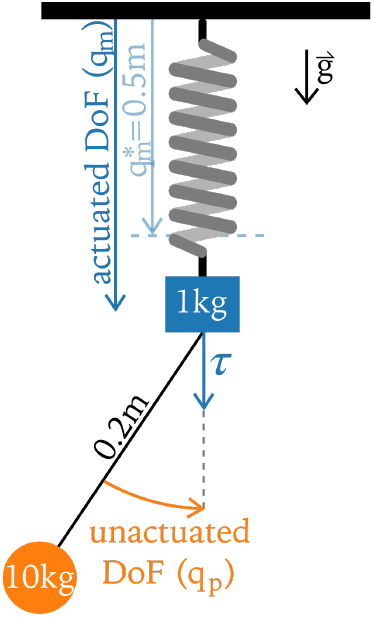
Spring-mass-pendulum model of Ex. III-C.

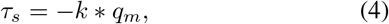

with *k* the spring stifness constant.

The OCP was composed of two phases each lasting for 5 s, with 50 shooting nodes. In the first phase, no objective function was minimized and *τ* was constrained to be 0, letting the mass oscillating freely. Then, in the second phase, a cost function (Eq. 5) was minimized, to enforce a reference position 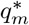 of the mass. This objective function, exclusively composed of Lagrange terms, was formulated as follows:

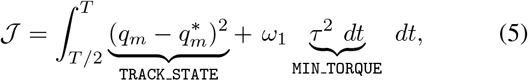

with 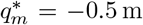 and *ω*_1_ = 10^*−6*^ and T is the duration of the movement. The first term of the objective function (Eq. 5) acts as a position controller for the mass. The second was added for control regularization.

During the first phase, the mass is passively oscillating around its stationary position due to the spring force (Fig. 7). At the beginning of the second phase, when an additional external force acts on the mass, it stabilizes around the targeted position. The standard deviation between the position and the targeted position is 0.04 m. This example highlights the possibility of using optimal control to find activation patterns compensating for external passive forces (e.g., ortheses flexibility, contact surface deformation, interaction between two models, etc.).

**Fig. 7:**
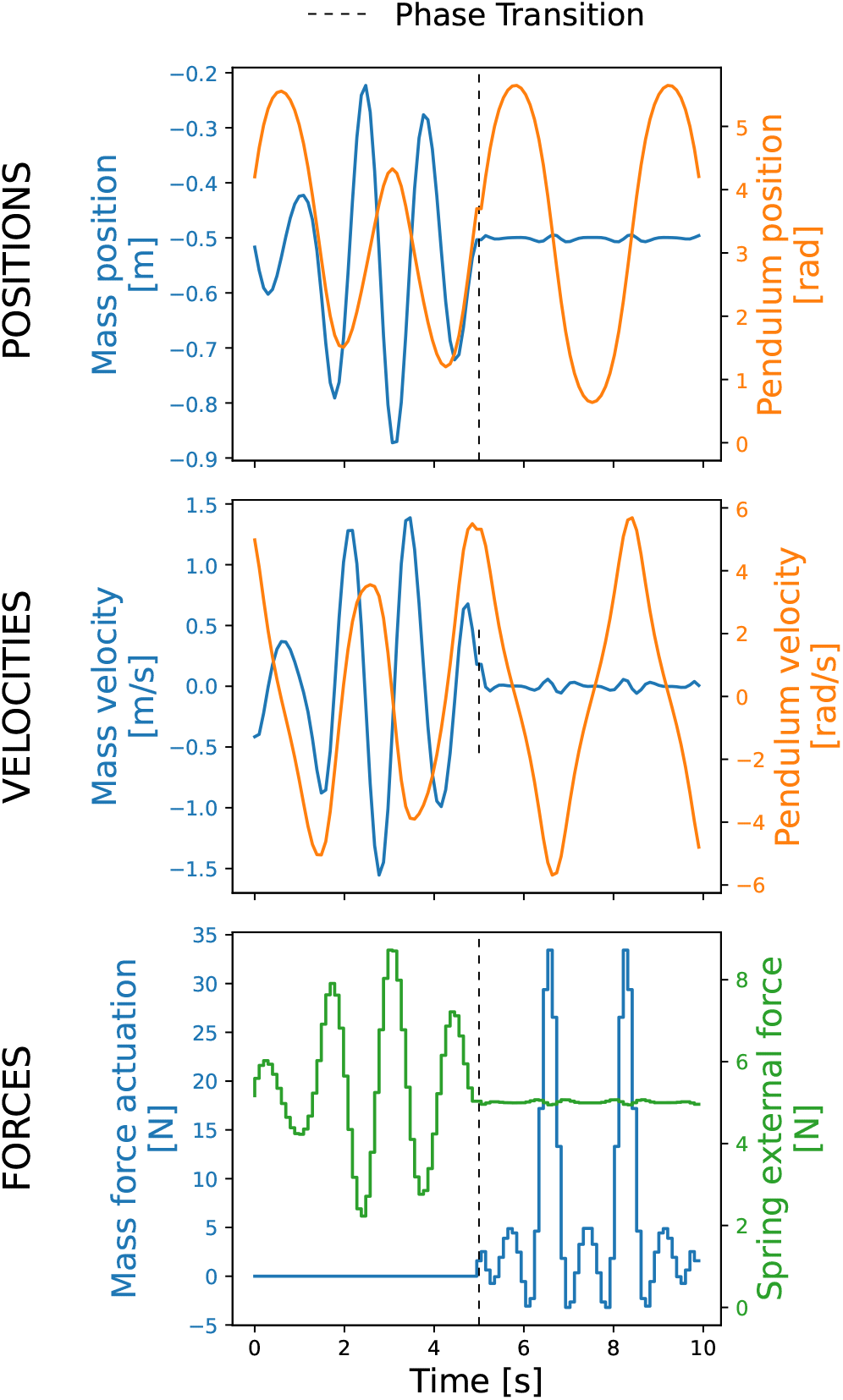
Two-phases kinematics of the mass-pendulum-spring system. Gray dashed lines show the phase transition, blue lines are related to the mass (position velocity and external force acting on it), red lines are related to the pendulum (position and velocity) and the green line depicts the spring force.

### D. Multiphase activation driven walking cycle

This example is presented to introduce *Bioptim*’s ability to deal with a multiphase locomotion estimation problem including muscle actuation and contact forces. The goal was to estimate muscle activations by tracking markers trajectories and ground reaction forces and moments. The model was a 3D leg with 12 DoFs (6-DoF pelvis, 3-DoF hip, 1-DoF knee and 2-DoF ankle), driven by 19 muscle activations and residual joint torques to compensate for potential muscle actuation weaknesses. The gait cycle was defined from the first heel strike to the end of the swing phase discretized into 90 shooting intervals. To approximate the natural rolling of the foot, the stance was divided into three phases (heel, flatfoot and forefoot contacts) of fixed duration deduced from experimental force platform data and markers position (0.05, 0.36 and 0.16 s). The swing phase lasted 0.38 s. The interaction between the ground and the foot was modeled using a four-contact points model located at the heel and the forefoot (first, fifth metatarsi and hallux). The optimization problem consisted in minimizing the errors between predicted (**m**) and reference (**m**^*^) markers trajectories, predicted (ℱ, ℳ) and reference (ℱ^**\***^, ℳ^*^) ground reaction forces and moments at all contact points. A regularization term on muscle activations (**a**) was also added (least-activations) as well as a penalization term on the residual torques (***τ***):

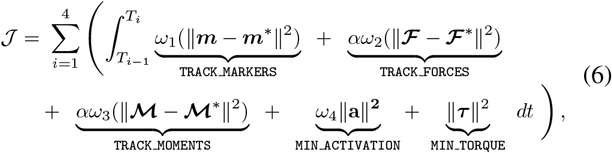

where *ω*_1_ = 10^5^, *ω*_2_ = *ω*_3_ = 10^*−1*^, *ω*_4_ = 10 are weighting factors, *T*_0_ = 0 and *T*_*i*_, *i ∈* [1, 2, 3, 4], are the final time of the i^*th*^ phase. Ground reaction forces and moments were only tracked during the stance phase, hence *α* = 0 during the swing phase and *α* = 1 otherwise. Non-slipping (`NON_SLIPPING`) and unilateral contact force (`CONTACT_FORCE`) constraints were added to prevent the foot from slipping and pulling from the ground. In between phases, the use of the `PhaseTransition.IMPACT` state transition allowed to represent the gain or loss of contact(s) in the dynamics (e.g., [27] swing phase to heel strike).

Tracking experimental data allowed to reproduce leg motion during the walking cycle (Fig. 8). The root mean square tracking error on markers trajectories was 27 mm (mean errors on pelvic and foot markers were 7.5 mm and 14.7 mm, respectively). Concerning ground reaction forces tracking, the root mean square error was 27 N. During the stance phase, Gluteal muscles and Vastus Medialis were mainly activated during the loading response (10 %) and hamstrings during initial contact (1 %) (Fig. 8). These results were similar to the characteristic average activity patterns of the lower limb muscles during locomotion described in [28]. The transition from stance to swing (60 % - 70 %) was highly actuated by hip flexors (Iliopsoas and Rectus Femoris) and leg muscles (Gastocnemius Medialis and Tibialis Anterior).

**Fig. 8:**
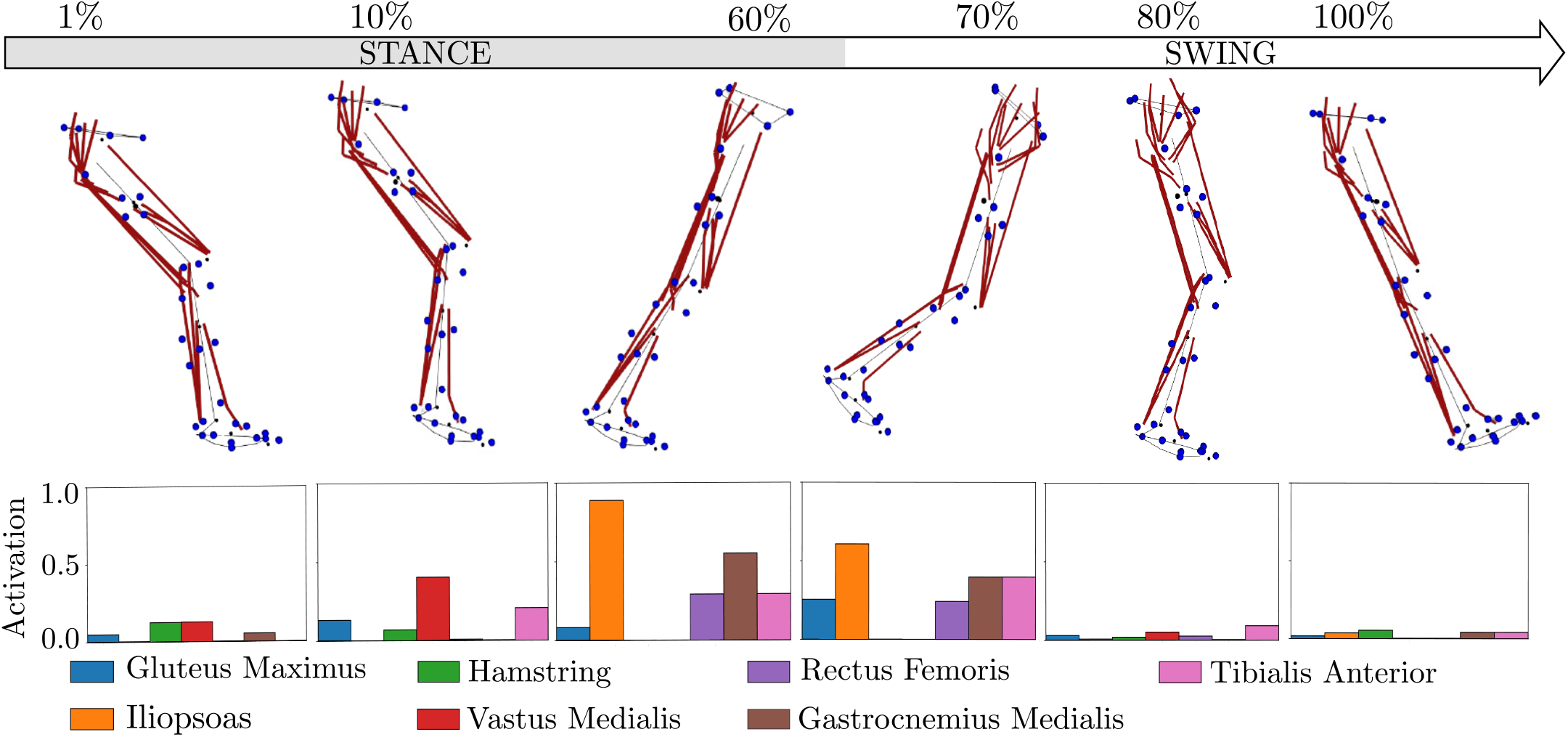
Snapshots of a walking gait cycle driven by muscles activation with histogram of muscle activations below. The red lines represent muscles lines of action and the blue points depict the tracked markers. The activation of the Gluteus Maximus is the mean of its three parts and the Hamstring is the mean activation of the Semimembranous, Semitendinous and Biceps Femoris.

### E. Moving Horizon Estimation of Shoulder Elevation

This example is presented to introduce *Bioptim*’s ability to provide real-time estimation of biomechanical variables. The goal was to perform a real-time estimation of dynamically consistent joint kinematics and muscle forces, using a moving horizon estimation (MHE) approach (i.e. an optimization approach that uses a series of measurements observed over time). A shoulder elevation motion was performed with a 4-DoF (**q**) arm actuated by 19 Hill-type muscle elements. The control inputs of the model were the muscle activations (**a**). The MHE implementation consists in splitting the OCP into a succession of smaller one for processing fixed-size subsets of the tracking data moving forward in time. Each time one subproblem is solved, a new measurement is added, the oldest one is discarded and a new subproblem is defined. Due to their similarities, the solution of the previous OCP is a good initial guess to the new one. The dynamical consistency of the final solution is enforced by continuity constraints on the initial state. Each objective function (Eq. 7) was written as the sum of three terms: tracking reference joint angles (**q**^*^), states and muscle activations regularizations (i.e., least-square criteria):

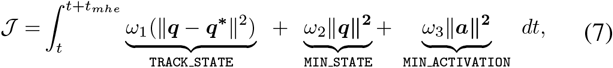

where *ω*_1_ = 10^3^, *ω*_2_ = 10, *ω*_3_ = 10^2^ and *t*_*mhe*_ is duration of each sub-problem.

In this example, reference data of an 8 s series of four arm elevations were generated at 100 Hz, by computer simulation. A centered Gaussian noise (mean = 0, std = 0.005 *q*^*^(*t*)) was added to *q*^*^, to simulate experimental-like joints angle measurements. Using a windows size of 7 nodes (i.e., 210 ms), the estimator ran at about 33 Hz (one in three reference data frame was sent to the estimator to simulate experimental-like conditions), i.e., two and half times faster than standard biofeedback (13 Hz, [29]). The MHE was able to forecast the movement kinematics with a root mean square error of 1.3 *±* 0.7^*°*^ while providing a realistic estimation of muscle forces close to the ground truth with a root mean square error of 11.1 *±* 14.9 N (Fig. 9).

**Fig. 9:**
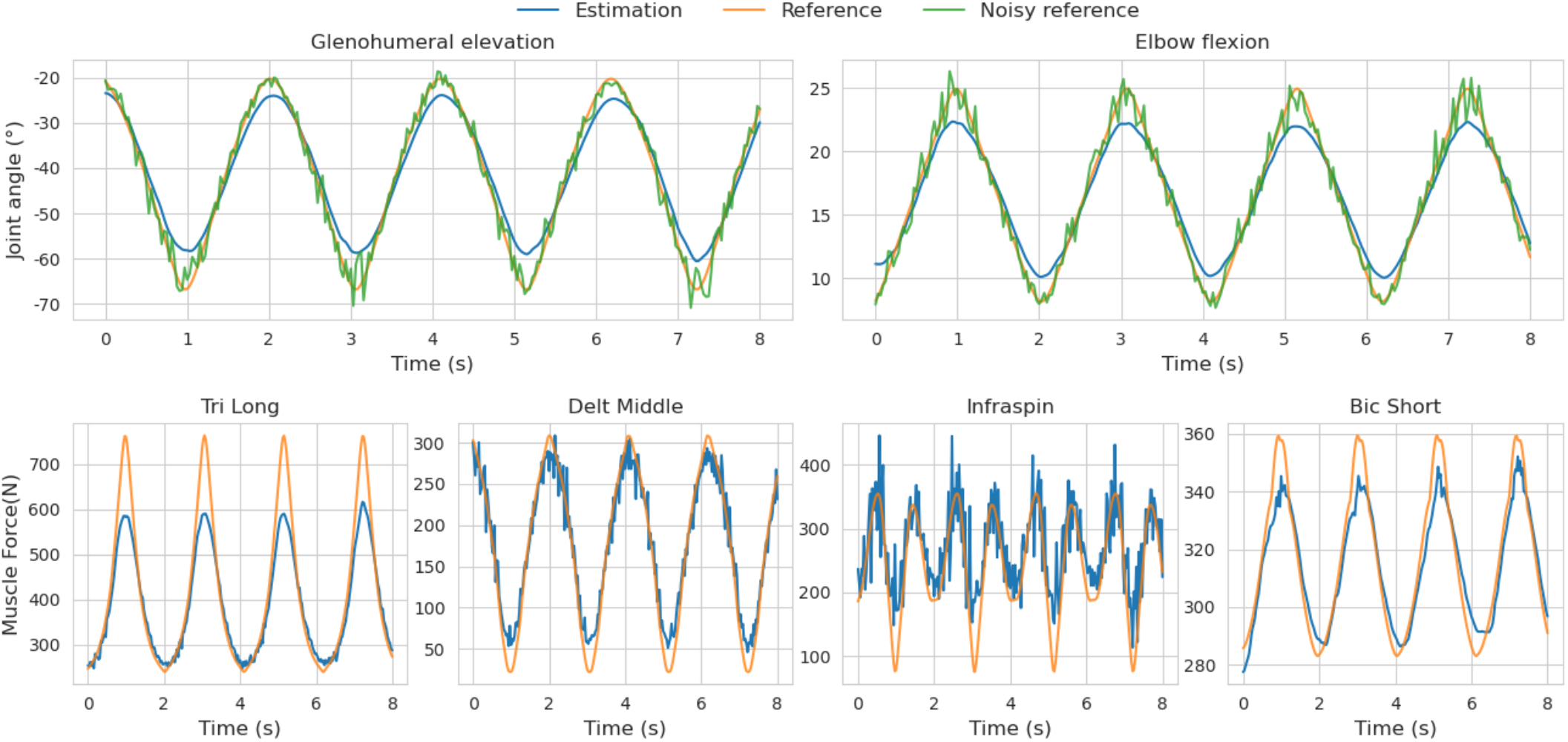
Ex. III-E. Top row - Real-time estimated joint angles (blue), ground-truth joint angles (orange) and tracked noisy joint angles (green) for a cyclic motion of the arm. Bottom row - Real-time estimated muscle forces (blue) and ground truth muscle forces (orange) for the same motion. Only four muscles with significative action (peak forces *>* 15 N), on the two selected DoFs, are shown. Muscle abbreviations stand for (from left to right): Triceps Long head, Deltoid Middle, Infraspinatus, Biceps Brachial Short head.

### F. Multiphase vertical jumper

This example was designed to introduce *Bioptim*’s ability to reduce the number of degrees-of-freedom (DoF) of a model via the `BiMapping` feature, to account for nonlinear boundaries on the controls, and to solve complex multiphase OCP. A total of five phases were used to describe the various dynamics of the jump, namely the push-off phase (i.e., flat foot (two floor contacts^2^) and then toe only (one contact)), flight (free fall, i.e., no contact) and landing (toe (one contact) and then flat foot (two contacts)). When a contact was added between two phases, we used the build-in inelastic collision phase transition `PhaseTransition.IMPACT` to compute the velocities of the kinematic chain at the beginning of the post-impact phase. A pseudo-2D full-body symmetrized model consisting of 3 DoFs at the pelvis (forward and upward translations, tranverse rotation), 1 DoF at the upper limb (shoulder flexion), and 3 DoFs at the lower limb (hip, knee and ankle flexion) was used. Since this is a full-body model, the root segment (i.e., the pelvis) was left uncontrolled, reducing the number of control variables to four, namely the shoulder, hip, knee and ankle flexions. The objective function with the most important weight was a Mayer objective computed at the end of the push-off phase consisting in maximizing the jump height (*h*) from the free fall equations applied to the center of mass. The remaining objective functions were regularization terms and terms that favoured a human-like solution.

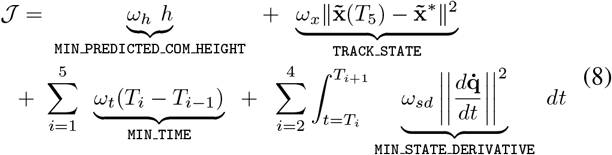

where *T*_*i*_ with *i ∈* [1, 2, 3, 4, 5] are the final times of the i^*th*^ phase respectively, and *T*_0_ = 0; *ω*_*h*_ = − 100 is the weight of the jump height term defined negative to maximize it; *ω*_*t*_ = 0.1, *ω*_*sd*_ = 0.1 and *ω*_*x*_ = 1.0 are the weights of their respective objective functions; 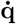 is the generalized velocities part of the state vector **x**; and 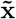 is the state vector excluding the translations of the root segment. The 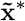 corresponds to a reference static position of the avatar with its knee slightly flexed and its arms horizontal.

Joint angles were bounded to human-like limits. The first node of the first phase was enforced to be equal to **x**^*^ (i.e., including the translations of the root segment to be at the origin). Joint velocities were arbitrarily bounded to human-like limits. Joint torques were bounded with nonlinear torque/angle/velocity relashionships measured on a high-level athlete using an isokinetic dynamometer [30]. Non slipping (`NON_SLIPPING`) and unilateral (`CONTACT_FORCE`) contact force constraints were added to prevent the contact points from slipping and pulling on the ground. During the ground phases, the heels had to remain over the floor. To speed-up the convergence with *Ipopt*, the problem was first solved using a BFGS Hessian approximation for 200 iterations. Then, starting from this first solution, the problem was re-optimized, with the exact-Hessian computations.

The optimized solution was obtained in 148 iterations of the exact-Hessian optimization resulting in a 1.28 m jump height. The optimized time for phases 1 to 5 were 0.70, 0.05, 0.99, 0.36, 0.21 s. The solution reproduced a human proximo-distal strategy (Fig 10), i.e., activating large segments first (for instance the torso) and sequentially adding more distal segments, consequently ending up with the feet.

**Fig. 10:**
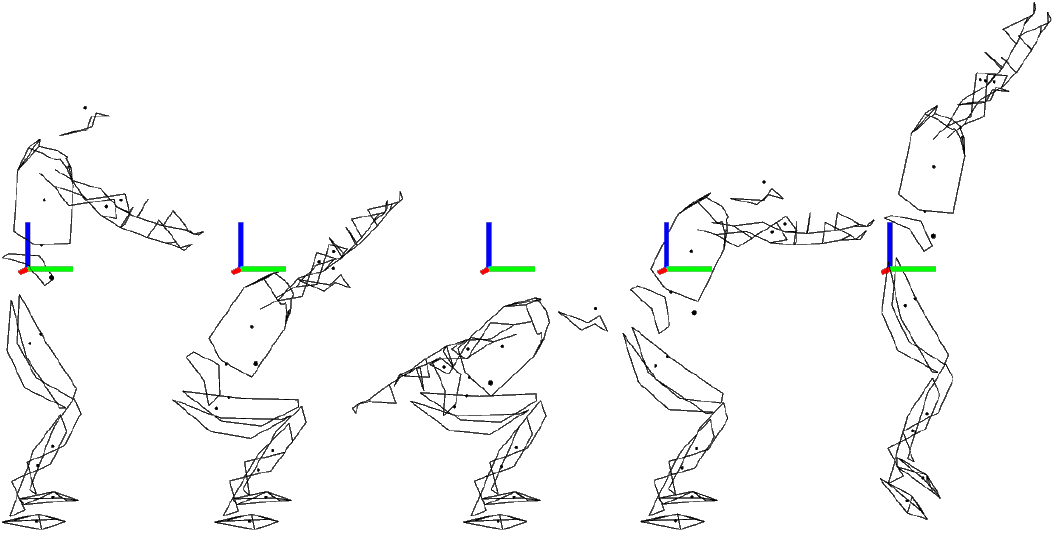
Snapshots of the push-off phase of a vertical jump (Ex. III-F). The avatar reproduces a human-like jump movement. The first four positions represent the first phase of the optimization (i.e., heel and toe in contact with the floor) and the fifth position depicts the end of the second phase (i.e., only the heel in contact with the floor)

## IV. Discussion

The purpose of *Bioptim* is to solve a variety of biomechanical OCPs with minimal user effort and high performances in terms of computational time. The main features illustrated by the six provided examples are (Tab. I):

- the possibility to use torque- or muscle-driven models (and their combinations);
- a variety of ready-to-use cost functions, constraints and dynamics (with and without contacts)…
- … easily customizable in Python when required by the user;
- the possibility to solve advanced OCPs (possibly multiphase) in a few seconds or minutes, that previously took us hours;
- the interface with two different NLP solvers

In addition, every feature of *Bioptim* is thoroughly illustrated by the examples of the getting_started folder (parameter optimization, custom objects, etc.). In the following, several aspects of *Bioptim* are discussed.

### A. Direct multiple shooting-based

While the debate remains about the performances of direct collocations versus direct multiple shooting [7], [3], the development of *Bioptim* was oriented toward the latter, because: *i)* it allows to select effortlessly an arbitrary accuracy for the integration (e.g., order and numbers of RK steps); *ii)* it allows to use multiple shooting-based fast NLP solvers such as *ACADOS*. Concerning the integration, either internally or via *ACADOS*, several schemes are implemented in *Bioptim* (RK4, RK8, implicit RK). While IRK showed better convergence in our experience with hard problems in *ACADOS*, RK4 showed to be a good speed/robustness tradeoff in most of the cases. In contrast to what is claimed in [3], direct multiple shooting is not a limitation to the performances (cost value and time to convergence), since, in our experience, the performances of *Bioptim* often outperform state-of-the-art results.

### B. Automatic differentiation

One of the reasons explaining the performances of *Bioptim* is the rewriting of the core software, *RBDL* [31] and *Biorbd* implementing the dynamics, into *CasADi* symbolics to automatically provide the exact Jacobians and Hessians of the resulting NLP. The gain in accuracy for the calculation of derivatives leads to shorter convergence times (due to much less iterations) and to optimal solutions reached with lower tolerances. This last aspect must be emphasized for complex motions (fast, highly dynamics ones), because, for instance when using *Ipopt*, an optimal solution obtained with a convergence criterion of 10^−2^ is very unlikely to be dynamically sound; i.e., it would diverge when forwardly integrating the controls in a single-shooting manner. A lower tolerance (10^−6^ or 10^−8^), which is only reachable with exact derivatives—for most of OCPs in biomechanics—, is expected to lead to better forward dynamics results.

### C. Python based, but fast!

*Bioptim* was thought as an interface, and was therefore written in Python to allow the user to easily combine existing cost functions or constraints and self-implemented ones, to switch from one solver to another, etc. We believe this feature to be of importance given that the biomechanics community is mainly composed of software users rather than developers. Therefore, providing a custom interface in Python rather than in C++, was a driving objective of our work to facilitate a rapid appropriation by the community. Since flexibility and ease-of-use should not compromise the performances, the integration is multi-threaded and all the inside computations are expressed as C++ *CasADi* graphs, interfaced with C++ NLP solvers. These graphs can either be built in `casadi.MX()` or `casadi.SX()`. The latter requires more RAM for building the problem but is faster to solve. While both may be used with *Ipopt, ACADOS* is only compatible with `casadi.SX()`. By leveraging the speed of `casadi.SX()` graphs, we were able to estimate muscle forces in real time using *ACADOS* on a standard laptop (Ex. III-E). For a more in-depth analysis of the real-time estimation capabilities of *Bioptim*, see [2].

Alongside with the 3D visualizer *Bioviz* that animates the solution, *Bioptim* proposes a series of online-generated figures, inspired by the real-time graphics from *Muscod-II* [24], [25], to visualize the optimized variables at each iteration of the solver. This is made with minimal computational cost thanks to the multiprocessing Python toolbox. Our implementation leverages the *Python pickle* library for easily saving and loading OCPs for, e.g., post-processing analysis. Finally, every layer (integration, optimization, visualization) of *Bioptim* is optimized to be flexible and fast.

### D. Fast vs robust NLP solvers

Fast solvers, such as *ACADOS*, offer the opportunity to use multi-start approaches on complex problems, to circumvent the obstacle of local minima [17], [11]. It also allows to get meaningful initial solutions from simpler problems, for guiding the resolution of the harder problems. On the other hand, robust solvers, such as *Ipopt*, are convenient when the user lacks information about the sought solutions and thus cannot guide the solver through a good initial guess. For biomechanics applications, the complementary characteristics of the interfaced solvers is a really useful tool. Moreover, *Bioptim*’s full compatibility with *CasADi* provides the opportunity to use any solver already interfaced with it, including third-party software such as *SNOPT, WORHP* [32] and *KNITRO* [33] (not tested yet).

### E. Multiphase

Biomechanics studies often face changing dynamics or objective functions due to the loss or gain of contacts or time-varying biomechanical tasks. When tracking such a motion or trying to predict it, these changes translate into multiphase OCP. This is one of the reported drawbacks of *OpenSim Moco*, which does not provide this feature yet. *Bioptim*, however, is able to handle multiphase OCPs, although they can currently only be solved with *Ipopt* (see Exs. III-D and III-F).

### F. From constraints to objectives: easy problem relaxation

As stated in Sec. II.B, there exists a correspondence between most of the pre-implemented `Constraints` and `Objective functions`. This is intended to allow for easy relaxation when the problem is reluctant to converge. For instance, when a biomechanical task requires the final configuration of the model to be enforced (reaching, cyclic motions, sports, etc.), one should first use a `Constraint` (e.g., `TRACK STATE`). If the convergence is challenging, just turning this constraint into its namesake Mayer `Objective function`, with a heavy weight, should help the solver.

### G. Limitations

*Bioptim* is already a mature solution for solving biomechanical OCP. However some limitations should be raised. First, it is based on *Biorbd* which is not as advanced as *OpenSim* or *AnyBody* (AnyBody Technology) in terms of biomechanical features and audience. Nevertheless, *Biorbd* is actively maintained, fast and *CasADi*-compatible for automatic differentiation. The variety of proposed examples highlighted simple to advanced models. Even if defining a new model was made straightforward thanks to the `.bioMod` file format, *biorbd* does not include a GUI for building models. Some Opensim models can be translated into `.bioMod` but *Biorbd* does not yet support multiple wrapping objects, non-orthogonal DoFs between bodies, compliant contact force models ([34]) or muscle-tendon equilibrium. As seen in [22], wrapping objects are rare due to the computational cost and required optimization when a line of action is in contact with more than one object, which compromises automatic differentiation. Via-points and pre-processed moment arms [35] (to be expressed as polynomial functions of crossed DoFs) are often preferred.

### H. Future directions

*Bioptim*, code name *PaperWork* (Version 1.1.0), was released in February 2021, with all the features presented in this communication. Some improvements are expected in a near future. First, a graphical model builder is planned in *Biorbd*, to easily generate .*bioMod* files. Also, models of muscular fatigue are to be included in *Bioptim*, to predict adapted motor strategies for long or demanding motions. The formulation of moving horizon schemes (MHE, Nonlinear Model Predictive Control) will be pre-implemented, with efficient warm-starting heuristics, to facilitate their use. The implementation of muscle-tendon equilibrium is planned for fast movements or those with large ranges of motions. It will require an additional optimization step to achieve the equilibrium as done in *CEINMS* [1] or the addition of muscle lengths as state variables, as in [35]. Moreover, an effort will be made to extend the compatibility of *ACADOS* with all the features of *Bioptim* (multiphase, nonlinear constraints, etc.). Finally, we plan to add an inverse optimal control module to *Bioptim* and muscle synergy dynamics to improve motion predictions [36].

## Acknowledgment

This study and the *Biorbd* library development was partly funded by a scholarship of the Vanier program (BM), the Canada First Research Excellence Fund via the TransMedTech Institute (FB) and the NSERC Discovey Programme (MB). *Bioptim* acts as a catalyst in our group and several students contributed to this library. Thank you to Théophile Gousselot, Paul Wegiel, Ariane Dang, Valentin Thiron and André Venne.

## Footnotes

1 [link](https://github.com/pyomeca/bioptim) – DOI: 10.5281/zenodo.4562883

2 A contact is defined as a point where forces are applied to cancel its acceleration.

## References

[1] C. Pizzolato, D. G. Lloyd, M. Sartori, E. Ceseracciu, T. F. Besier, B. J. Fregly, and M. Reggiani, “Ceinms: a toolbox to investigate the influence of different neural control solutions on the prediction of muscle excitation and joint moments during dynamic motor tasks,” Journal of Biomechanics, vol. 48, no. 14, pp. 3929–3936, 2015.

[2] F. Bailly, A. Ceglia, B. Michaud, D. M. Rouleau, and M. Begon, “Real-time and dynamically consistent estimation of muscle forces using a moving horizon emg-marker tracking algorithm—application to upper limb biomechanics,” Frontiers in Bioengineering and Biotechnology, vol. 9, p. 112, 2021. [Online]. Available: https://www.frontiersin.org/article/10.3389/fbioe.2021.642742

[3] S. Porsa, Y.-C. Lin, and M. G. Pandy, “Direct methods for predicting movement biomechanics based upon optimal control theory with implementation in opensim,” Annals of Biomedical Engineering, vol. 44, no. 8, pp. 2542–2557, 2016.

[4] L. Modenese, “Awesome biomechanics,” https://git.io/JtdLh, 2020.

[5] M. Febrer-Nafr’ sia, R. Pallarès-Lo’pez, B. J. Fregly, and J. M. Font-Llagunes, “Comparison of different optimal control formulations for generating dynamically consistent crutch walking simulations using a torque-driven model,” Mechanism and Machine Theory, vol. 154, p. 104031, Dec. 2020.

[6] M. Ezati, P. Brown, B. Ghannadi, and J. McPhee, “Comparison of direct collocation optimal control to trajectory optimization for parameter identification of an ellipsoidal foot–ground contact model,” Multibody System Dynamics, vol. 49, no. 1, pp. 71–93, May 2020.

[7] M. Diehl, H. G. Bock, H. Diedam, and P.-B. Wieber, “Fast direct multiple shooting algorithms for optimal robot control,” in Fast Motions in Biomechanics and Robotics. Springer, 2006, pp. 65–93.

[8] J. Koschorreck and K. Mombaur, “Modeling and optimal control of human platform diving with somersaults and twists,” Optimization and Engineering, vol. 13, no. 1, pp. 29–56, 2012.

[9] M. Felis and K. Mombaur, “Modeling and optimization of human walking,” in Modeling, Simulation and Optimization of Bipedal Walking. Springer, 2013, pp. 31–42.

[10] E. Charbonneau, F. Bailly, L. Danès, and M. Begon, “Optimal control as a tool for innovation in aerial twisting on a trampoline,” Applied Sciences, vol. 10, no. 23, p. 8363, 2020.

[11] F. Bailly, E. Charbonneau, L. Danès, and M. Begon, “Optimal 3d arm strategies for maximizing twist rotation during somersault of a rigid-body model,” Multibody System Dynamics, pp. 1–17, 2020.

[12] M. Giftthaler, M. Neunert, M. Stäuble, and J. Buchli, “The control toolbox—an open-source c++ library for robotics, optimal and model predictive control,” in 2018 IEEE International Conference on Simulation, Modeling, and Programming for Autonomous Robots (SIMPAR). IEEE, 2018, pp. 123–129.

[13] F. Bailly, J. Carpentier, B. Pinet, P. Souères, and B. Watier, “A mechanical descriptor of human locomotion and its application to multi-contact walking in humanoids,” in 2018 7th IEEE International Conference on Biomedical Robotics and Biomechatronics (Biorob). IEEE, 2018, pp. 350–356.

[14] M. R. Yeadon and M. J. Hiley, “The mechanics of the backward giant circle on the high bar,” Human Movement Science, vol. 19, no. 2, pp. 153–173, 2000.

[15] M. Begon, M. J. Hiley, and M. R. Yeadon, “Effect of hip flexibility on optimal stalder performances on high bar,” Computer Methods in Biomechanics and Biomedical Engineering, vol. 12, no. 5, pp. 575–583, 2009.

[16] F. Leboeuf, G. Bessonnet, P. Seguin, and P. Lacouture, “Energetic versus sthenic optimality criteria for gymnastic movement synthesis,” Multibody System Dynamics, vol. 16, no. 3, pp. 213–236, 2006.

[17] A. Huchez, D. Haering, P. Holvoët, F. Barbier, and M. Begon, “Local versus global optimal sports techniques in a group of athletes,” Computer Methods in Biomechanics and Biomedical Engineering, vol. 18, no. 8, pp. 829–838, 2015.

[18] A. Wächter and L. T. Biegler, “On the implementation of an interior-point filter line-search algorithm for large-scale nonlinear programming,” Mathematical Programming, vol. 106, no. 1, pp. 25–57, 2006.

[19] P. E. Gill, W. Murray, and M. A. Saunders, “Snopt: an sqp algorithm for large-scale constrained optimization,” SIAM Review, vol. 47, no. 1, pp. 99–131, 2005.

[20] R. Verschueren, G. Frison, D. Kouzoupis, N. van Duijkeren, A. Zanelli, R. Quirynen, and M. Diehl, “Towards a modular software package for embedded optimization,” IFAC-PapersOnLine, vol. 51, no. 20, pp. 374–380, 2018.

[21] J. A. Andersson, J. Gillis, G. Horn, J. B. Rawlings, and M. Diehl, “Casadi: a software framework for nonlinear optimization and optimal control,” Mathematical Programming Computation, vol. 11, no. 1, pp. 1–36, 2019.

[22] C. L. Dembia, N. A. Bianco, A. Falisse, J. L. Hicks, and S. L. Delp, “Opensim moco: musculoskeletal optimal control,” PLOS Computational Biology, vol. 16, no. 12, p. e1008493, 2020.

[23] M. A. Patterson and A. V. Rao, “Gpops-ii: a matlab software for solving multiple-phase optimal control problems using hp-adaptive gaussian quadrature collocation methods and sparse nonlinear programming,” ACM Transactions on Mathematical Software (TOMS), vol. 41, no. 1, pp. 1–37, 2014.

[24] D. B. Leineweber, I. Bauer, H. G. Bock, and J. P. Schlöder, “An efficient multiple shooting based reduced sqp strategy for large-scale dynamic process optimization. part 1: theoretical aspects,” Computers & Chemical Engineering, vol. 27, no. 2, pp. 157–166, 2003.

[25] D. B. Leineweber, A. Schäfer, H. G. Bock, and J. P. Schlöder, “An efficient multiple shooting based reduced sqp strategy for large-scale dynamic process optimization: Part ii: software aspects and applications,” Computers & Chemical Engineering, vol. 27, no. 2, pp. 167–174, 2003.

[26] B. Houska, H. J. Ferreau, and M. Diehl, “Acado toolkit—an open-source framework for automatic control and dynamic optimization,” Optimal Control Applications and Methods, vol. 32, no. 3, pp. 298–312, 2011.

[27] M. L. Felis and K. Mombaur, “Synthesis of full-body 3-d human gait using optimal control methods,” in 2016 IEEE International Conference on Robotics and Automation (ICRA), 2016, pp. 1560–1566.

[28] D. A. Winter, Biomechanics and motor control of human gait: normal, elderly and pathological - 2nd edition. University of Waterloo Press, 1991, vol. Ed2. [Online]. Available: https://trid.trb.org/view/770965

[29] O. A. Kannape and O. Blanke, “Self in motion: sensorimotor and cognitive mechanisms in gait agency,” Journal of Neurophysiology, vol. 110, no. 8, pp. 1837–1847, 2013.

[30] M. I. Jackson, “The mechanics of the table contact phase of gymnastics vaulting,” Jan. 2010.

[31] M. L. Felis, “Rbdl: an efficient rigid-body dynamics library using recursive algorithms,” Autonomous Robots, pp. 1–17, 2016. [Online]. Available: http://dx.doi.org/10.1007/s10514-016-9574-0

[32] D. Wassel, “Exploring novel designs of nlp solvers: architecture and implementation of worhp,” Ph.D. dissertation, Universität Bremen, 2013.

[33] J. Nocedal, “Knitro: an integrated package for nonlinear optimization,” in Large-Scale Nonlinear Optimization. Springer, 2006, pp. 35–60.

[34] G. Serrancol’ si, A. Falisse, C. Dembia, J. Vantilt, K. Tanghe, D. Lefeber Jonkers, J. De Schutter, and F. De Groote, “Subject-exoskeleton contact model calibration leads to accurate interaction force predictions,” IEEE Transactions on Neural Systems and Rehabilitation Engineering, vol. 27, no. 8, pp. 1597–1605, 2019.

[35] A. J. Van Den Bogert, D. Blana, and D. Heinrich, “Implicit methods for efficient musculoskeletal simulation and optimal control,” Procedia Iutam, vol. 2, pp. 297–316, 2011.

[36] J. P. Walter, A. L. Kinney, S. A. Banks, D. D. D’Lima, T. F. Besier, D. G. Lloyd, and B. J. Fregly, “Muscle synergies may improve optimization prediction of knee contact forces during walking,” Journal of biomechanical engineering, vol. 136, no. 2, 2014.

